# Tool skill impacts the archaeological evidence across technological primates

**DOI:** 10.1101/2024.06.10.598310

**Authors:** Lydia V. Luncz, Nora E. Slania, Katarina Almeida-Warren, Susana Carvalho, Tiago Falótico, Suchinda Malaivijitnond, Adrián Arroyo, Ignacio de la Torre, Tomos Proffitt

## Abstract

The archaeological record offers insights into our evolutionary past by revealing ancient behaviour through stone and fossil remains. Percussive foraging is suggested to be particularly relevant for the emergence of tool-use in our lineage, yet early hominin percussive behaviours remain largely understudied compared to flaked technology. Stone tool-use of extant primates allows the simultaneous investigation of their artefacts and the associated behaviours. This is important for understanding the development of tool surface modification, and crucial for interpreting damage patterns in the archaeological record. Here, we compare the behaviour and the resulting material record across stone tool-using primates. We investigate the relationship of nut-cracking technique and stone tool modification across chimpanzees, capuchins, and long-tailed macaques by conducting standardized field experiments with comparable raw materials. We show that different techniques likely emerged in response to diverse nut hardness, leading to variation in foraging success across species. Our experiments further demonstrate a correlation between techniques and the intensity of visible percussive damage on the tools. Tools used with more precision and efficiency as demonstrated by macaques, show fewer use wear traces. This suggests that some percussive techniques may be less readily identified in the archaeological record.

## 1. Introduction

While tool-use is not a unique characteristic of our species, our technologies and material culture are unparalleled in the animal world and are seen as a central aspect of being human. The evolutionary origin of our material culture has been reconstructed by investigating the lithic record of early hominins ^1^. The onset of tool-use is predominantly described through sharp-edged stone flakes with the earliest evidence potentially dating back to 3.3 million years ago ^2,3^. The characteristics of flakes are generally thought to be indicative of the deliberate behaviour that produced them and stone flaking is often considered a complex behaviour that requires advanced cognitive abilities such as the understanding of raw material properties, planning and advanced motor skills ^4,5^. However, some have suggested that stone flaking does not represent the first step in the evolutionary history of tool-use ^6^. Percussive activities may have predated flaking and therefore played a key role for the origins of hominin tool-use ^7–12^. Various percussive artefacts are known from Early Stone Age archaeological sites. These have been associated to both flaking activities such as passive hammer ^4^ and bipolar knapping ^13–15^, as well as extractive foraging activities such as plant processing ^16,17^ and bone breaking ^18^. While pounding implements have been identified in the hominin archaeological record, various percussive activities may remain unidentified due to the previous lack of diagnostic tools to reliably interpret their archaeological signature ^14^.

An increasing referential database of percussive use wear through experimental and non-human primate studies show that different pounding behaviours often result in variation of percussive damage ^19–22^. Furthermore, GIS analyses have been used to describe and quantify damage patterns on percussive elements, including both hammerstones and anvils ^23–26^. Whilst controlled experiments offer a means to quantify percussive use wear and to classify tools accordingly, interpretations of percussive artefacts can be substantially improved when the relationships between behaviour and damage patterns are well understood ^21^. Whilst early hominin behaviors are inferred and reconstructed from analysis of a fragmentary archaeological record, living primates can function as a model species to further elucidate the potential behavioral diversity of early hominins ^8,27^.

Four non-human primate genera – chimpanzees (*Pan troglodytes verus*), three capuchin species (*Sapajus libidinosus, Sapajus xanthosternos, Cebus capucinus*), and long-tailed macaques (*Macaca fascicularis*) – engage in stone-tool-use. Amongst an array of stone tool behaviours specific for individual species, all these primates use stone tools to crack open nuts ^28–30^. Such a behaviour has also been suggested as a possible foraging activity of Plio-Pleistocene hominins ^13,16^. Foraging with tools offers a means to access otherwise unobtainable or costly food resources ^31–33^. Individual and species-specific skill thereby determine the relative benefit gained by engaging in tool behaviour. Benefits directly relate to the effort invested in tool-use. The higher the energetic intake relative to the energetic cost per unit of time, the higher the benefit ((energetic intake – energetic cost)/time), where more efficient tool-use yields greater benefits ^33,34^. For competitive foragers, quick and efficient nutrient intake is important ^34^. Efficiency of percussive foraging activities therefore has been investigated by assessing the caloric intake over time (nuts consumed per unit time), and the effort invested per food item consumed (hits per nut) ^34–37^. Various studies have shown that chimpanzees and capuchin monkeys adjust tool size and raw material to specific tasks to enhance their efficiency ^36–45^. The data on macaque percussive stone tool-use, albeit limited, indicate that they too adjust tool selection to a certain task ^25,46– 48^.

Proficient nut-cracking follows an operational sequence where the nut is placed on an anvil and hit with a hammer. A lack of precision could result in mishits or less effective strikes which would lower efficiency ^34,40^. At the same time, low levels of precision, when the hammerstone comes in contact with the anvil, result in larger areas of damaged stone surface. Stone-on-stone contact through partly or complete mishits of the nut leads to battered areas across the active surfaces ^21^. For instance, Arroyo and colleagues ^21^ describe direct impact on the anvil not only as a result of failings to strike the nut (mishits), but of strikes that push the nut off the active surface. These imprecise hits can lead to unsuccessful cracking attempts if the nut is lost or prolong the foraging process if the nut is being retrieved.

Here we compare nut-cracking behaviour of three stone tool-using primates (chimpanzees, bearded capuchins (*Sapajus libidinosus*) and long-tailed macaques) in their natural environment. Differences in cognition, morphological characteristics and foraging environments presumably affected how the behaviour emerged and developed through convergent evolution ^49^. This becomes especially apparent with the different nut-cracking techniques displayed across the three species. While chimpanzees and macaques crack nuts in a sitting position, often using one hand to lift the hammer a short distance (22; see Supplementary Video 1), capuchins usually employ a bipedal stance and use both hands to raise the hammer to face level ^49,50^. We expect these behavioural differences to be reflected in species nut cracking precision and therefore in the resulting damage patterns visible on their percussive tools.

To draw a meaningful comparison between species we set up controlled field experiments where each species was provided with the same hammer and anvil raw materials that are unavailable in their natural environment. These materials were imported from archaeological sites in eastern Africa to specify what percussive traces would look like on material available to early hominins. We asses species-specific skill by measuring efficiency and precision. Comparative values of efficiency are obtained via three measures: foraging duration (time needed to crack a nut), cracking efficiency (hits needed to crack a nut), and success rates (whether or not a selected nut was consumed after cracking attempts). Precision is assessed as the number of times a nut is pushed off the working surface in relation to the total number of hits per single nut. Comparative analysis of used stones is aimed at investigating differences in damage patterns on the tool material (hammerstones and anvils) across species. To assess the underlying reasons for observed diversity on nut cracking technique and success we experimentally test the nut structural resistance across environments. We predict that different techniques of tool handling affect the precision of hitting the nut. This will influence the efficiency (effort needed to crack open a nut) which in return affects the overall foraging success of a tool user. Taken those variables together we refer to this as skill. This composite measure of “skill” allows us to draw behavioral comparisons between species and to relate behavioral differences to tool surface modification.

## Results

### Comparisons of nut cracking behaviour across species

When cracking nuts, capuchins took a bipedal position and used both hands to raise the hammer (94%). In line with previous research ^38,51^, chimpanzees in this study cracked nuts in a sitting position (98%) always using one hand to hold the hammerstone. Similar to chimpanzees, macaques consistently took a sitting position (99.4%). Furthermore, macaques used their free hand to engage in shielding for the majority of events (62%) by positioning the hand so as to block the nut from potentially rolling or flying off the anvil. All three species showed no or low levels of grip adjustment, but capuchins adjusted their hammer stone handling most frequently (once per 2 nut cracking sequences) compared to chimpanzees (never) and macaques (once per 20 nut cracking sequences). While chimpanzees in a sitting position raised the hammerstone to a maximum of chest height in the majority of all events (71.43%), capuchins in bipedal stance raised the hammer to a maximum height of their forehead (48.71%), shoulder (25.18%) and overhead height (19.3%). Macaques raised their hammers to a maximum height of their chest (49%) and shoulder (47.5%) with roughly equal frequency. Furthermore, capuchins skipped feeding on successfully cracked nuts almost one in four times (22.5% of events), while chimpanzees and macaques failed to consume 9.5% and 1.6% of opened nuts respectively (see supplementary Table 10).

### Comparison of nut structural resistance

To assess nut structural resistance we conducted a standardized experiment across locally available nuts within the home range of our study animals. We measured how many times a standardized weight had to be dropped onto a nut from a standardized height in order for the nut to break open. This test showed that palm nuts used for capuchins were generally harder to crack than the oil palm nuts given to chimpanzees and macaques. Capuchin nuts needed 19.6 hits on average to open (min=1, max=78, median=14, n=101), compared to chimpanzee and macaque nuts which needed 4 (min=1, max=88, median=1, n=100) and 2.2 (min=1, max=7, median=2, n=100) hits on average. Nuts available to capuchins needed almost 10 times the amount of hits to crack open than the nuts available to chimpanzees and macaques. A one-way ANOVA in combination with a TukeyHSD test revealed a significant difference between nuts provided to capuchins and those used for chimpanzees and macaques (ANOVA: df=2, F=69.7, p<0.000; TukeyHSD: capuchins-chimpanzees p<0.000, capuchins-macaques p<0.000, chimpanzee-macaques p=0.5). This difference will affect efficiency and associated damage patterns on tools, which we consider throughout further interpretation of the data.

### Comparisons of foraging duration: (time required to consume nuts)

Overall, species differed in the time needed to consume a single nut (likelihood ratio test: X^2^ = 16.85, df = 2, *p* = 0.0002). Macaques generally required less time (mean = 7.33s, sd = 4.79s) to crack a nut than chimpanzees (mean = 32.26s, sd = 27,91s) and capuchins (mean = 19.23s, sd = 11.26s) (Table 2; see Supplementary Table 11 for model estimates and confidence intervals). Overall, hammer size did not have a significant impact on their foraging success (likelihood ratio test: X^2^ = 1.39, df=2; Table 2, see Supplementary Table 12 for model output of random effects). Fixed effects of the model explain 38.2% (marginal R2 = 0.382) of variance and the whole model can account for 61.8% (conditional R2 = 0.618) of variance within the response.

### Comparisons of cracking efficiency: (number of hits per nut)

The number of hits required to crack a nut varied between species (likelihood ratio test: X^2^ = 25.94, df = 2, *p*<0.0001). Macaques employed the fewest hits to open a nut with 1.3 mean hits (sd = 0.6), compared to capuchins (mean = 1.77, sd = 1.29) and chimpanzees (mean = 7.11, sd = 4.6) who use the highest number of hits (Table 2; see Supplementary Table 13 for model estimates and confidence limits). Hammer size affected the number of hits per nut (X^2^ = 6.5, df = 2, p=0.039): On average individuals needed 1.52 hits with the small hammer, 2.27 hits with the medium hammer and 1.94 hits with the large hammer (See Supplementary Table 14 for model output on random effects). The fixed effects part of the model explains 31.3% (trigamma marginal R2 = 0.313) of variance, while fixed and random effects together explain 41.2% (trigamma conditional R2 = 0.412) of variance within the response.

### Comparisons of overall success rates of processing naturally available nuts

Overall, species clearly differed regarding how successfully they cracked nuts (likelihood ratio test: X^2^ = 17.0, df = 2, p = 0.0002; see Supplementary Table 15 for model estimates and confidence intervals). Macaques successfully consumed 94% of nuts they attempted to crack, while chimpanzees and capuchins only foraged on 62% and 45% of nuts they attempted to crack respectively (Figure 1; Table 2). Hammer size did not affect cracking success (X^2^ = 1.76, df = 2, *p* = 0.414; see Supplementary Table 16 for model output on random effects). Fixed effects of the model account for 29.2% of variance (delta marginal R2 = 0.292) and the whole model explains 40.1% (delta conditional R2 = 0.401) of variance within the response.

**Figure 1:**
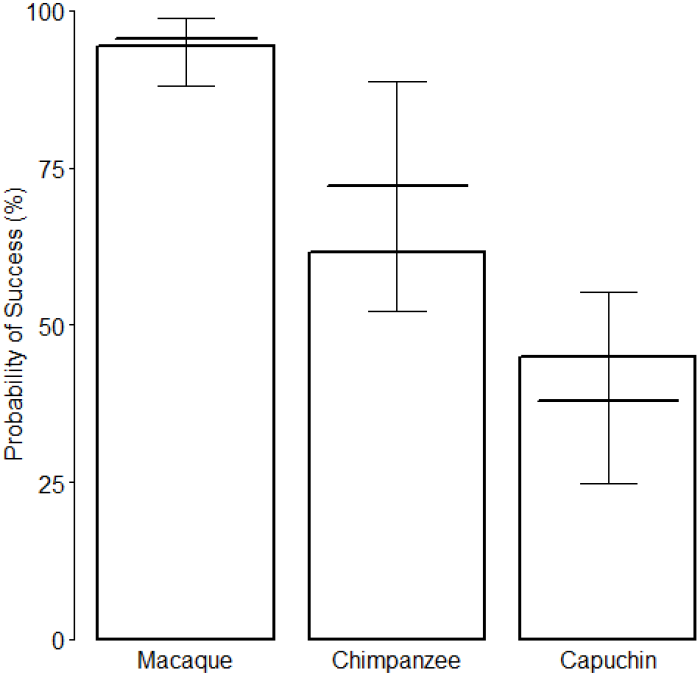
Success rates of nut cracking by three different stone tool using primate species. The y-axis depicts proportion of successful and unsuccessful cracking events in %. Horizontal lines with error bars show the fitted model and its 95% confidence interval for the fixed effect variable “Hammer Size” being at its average.

**Figure 2:**
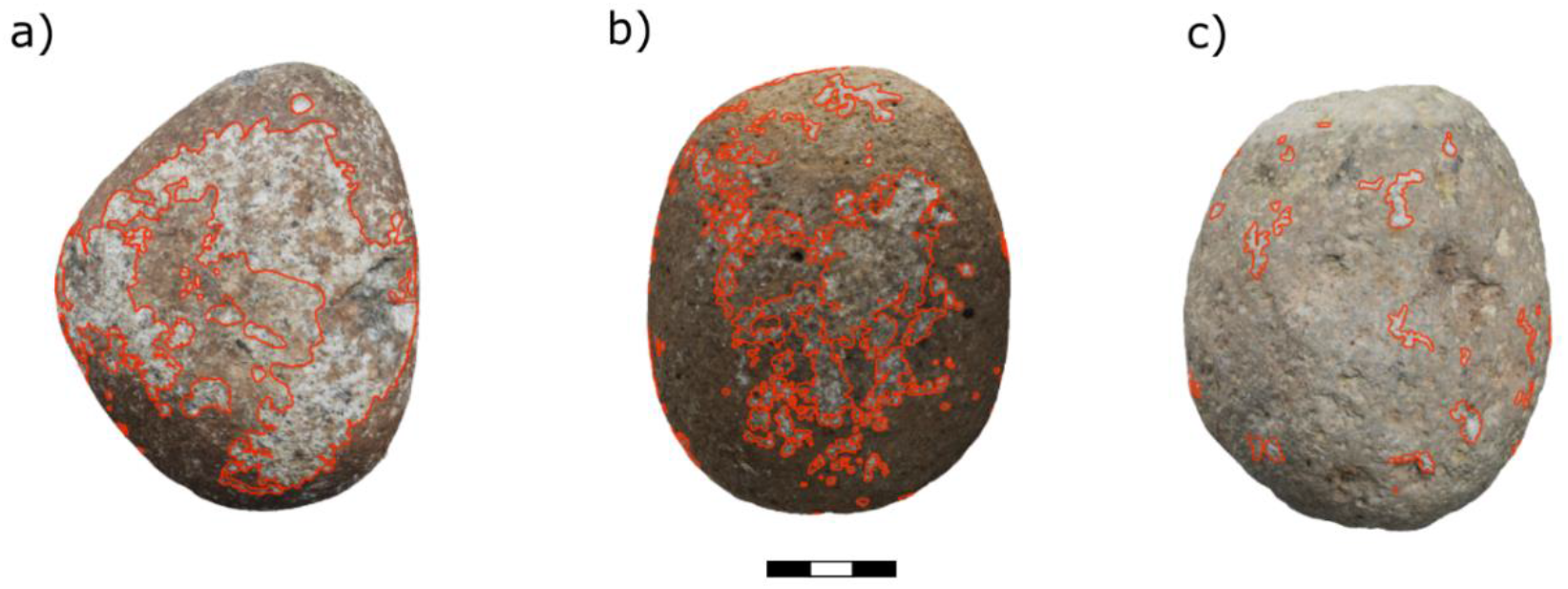
Example of damage patterns across species on hammerstones: Large hammerstones of (a) capuchins (b) chimpanzees and (c) macaques with damage outlined in red. (For all hammerstones see Supplementary Figure 1-3). Scale: 0.5cm per segment.

**Figure 3:**
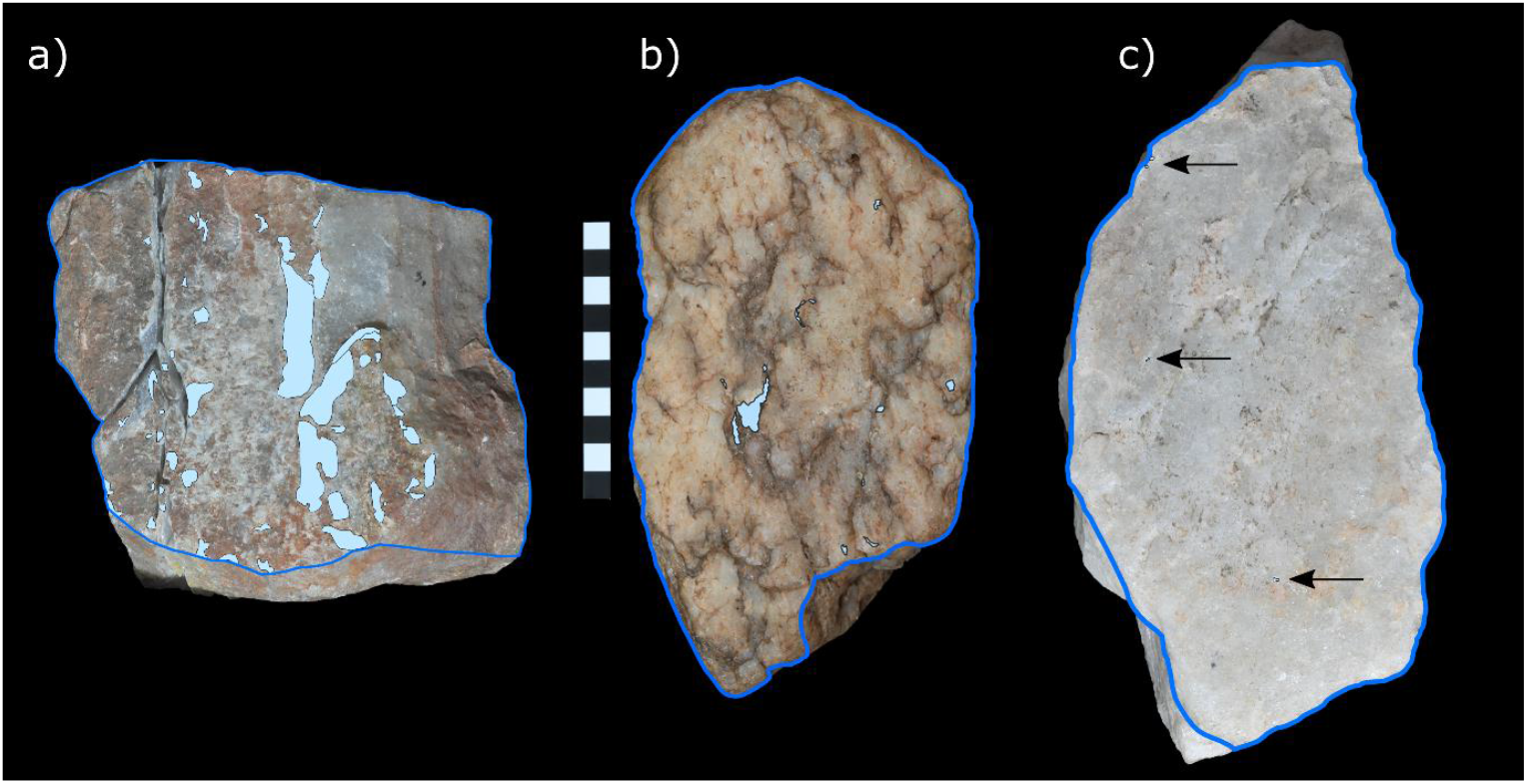
Percussive damage on anvils: Damage patterns across the active plane (outlined in blue) of each anvil used by (a) capuchins, (b) chimpanzees and (c) macaques. Note the almost complete absence of percussive damage on the macaque anvil. Scale = 10cm. Arrows point at single impact points.

### Comparison of striking precision

Species explains a significant amount of variance in the model (likelihood ratio test: X^2^ = 14.85, df = 2, p = 0.0006). Macaques are least likely to lose the nut during the cracking attempt (nut flies off per hit, mean = 0.08, sd = 0.24), followed by chimpanzees (nut flies off per hit, mean = 0.24, sd = 0.29) and capuchins (nut flies off per hit, mean= 0.40, sd = 0.45). For model estimates and confidence intervals see Supplementary Table 17. Hammer size does not affect how often the nut is being pushed off the working surface (X^2^ = 0.54, df = 2, p = 0.76). For model output on random effects see Supplementary Table 18 and 19. Fixed effects explain 17.8% (delta marginal R2 = 0.178) and the whole model explains 27.9% (delta conditional R2 = 0.279) of variance within the response.

### Comparisons of damage distribution patterns on hammerstones

Overall, there is a substantial difference between the damage pattern on the hammerstones and anvils seen across the three species. Capuchin hammerstones possess the highest percentage of area covered in macro percussive damage (mean PA = 15.5%). This is likely a result of the difference in nut hardness and therefore will be excluded from direct comparisons across species. Chimpanzee hammerstones exhibit a mean PA of 9% and macaque hammerstones possess the lowest mean PA value (1.8%). For detailed description of all variables for each hammerstone see Table 20 and Supplementary Figure 1-3. Damage on macaque hammerstones is limited to a few (mean = 67, range 6-53) medium-sized (mean area = 34.74mm^2^) areas on the tool (mean ED = 0.022). In contrast, damage on chimpanzee hammerstones is small (mean area = 10.77mm^2^) and widely distributed over the active surface (mean density=0.0090, n=524). Damage on capuchin hammerstones is centred on several (mean = 64.67, range 33-94), large (mean = 59.58mm^2^) areas on the hammerstones (mean ED = 0.133). Capuchins also show the single largest use wear area (LUW = 7.5%). For a detailed summary of the results, see Supplementary Table 20.

### Comparisons of damage distribution patterns on anvils

The damage pattern on the anvils generally follows the same species level variation observed for the hammerstones. The anvil used by capuchins possesses the highest number of individual use wear areas (n = 52), the greatest relative area (PA) of percussive damage (46.96%) as well as the largest single use wear area (LUW = 37.63%). As the effect of nut hardness cannot be separated from the damage pattern we also exclude the capuchin anvil from direct comparisons below. The anvil used by chimpanzees possess 12 individual areas of percussive damage, a PA of 0.72% and a LUW of 0.42%. The anvil used by macaques show the lowest degree of percussive damage with only 4 individual areas of damage, a PA of 0.02% and an LUW of 0.009% of the total surface area of the active plane (Summary of the results, Supplementary Table 21).

## Discussion

Extant primate stone tool-use offers an opportunity to investigate behaviours in direct combination with tool damage. This allows a greater understanding of how percussive use wear develops in relation to specific behaviours and techniques but in some cases also capabilities, such as skill. In this study, we compared tool behaviour of three stone-tool using primates and resulting damage patterns on stone material used for nut cracking.

Overall, capuchins have particular low success rates, consuming less than half of the nuts they attempt to crack, which results in fewer total feeding events. Further, capuchins show the lowest precision values with nuts flying off/being pushed off the anvil almost every second hit, roughly twice as often as for chimpanzees. Differences in both efficiency and precision likely relate to the technique applied to nut cracking resulting from the increase of nut-resistance. The naturally available nut for capuchins was about 10 times harder compared to what was available to macaques and chimpanzees. This requires substantial greater force per strike to crack pen a nut. Capuchins stand on their hind legs and raise the hammers above their heads when cracking nuts whereas the other primates usually sit down and do not raise their pounding tools higher than their mid waist (chimpanzees) or shoulder/chest (macaques). This can account for faster striking techniques of macaques and chimpanzees and further indicates that capuchins apply a lot more force onto a nut per hit. Greater force per hit can explain the relatively low numbers of strikes capuchins employ per nut. Performing strikes with greater force likely reduces an individual’s control over the tool and therefore precision of the strike^34,40^. Behavioural skill levels are thus likely to be dictated by the available resources within the natural environment of a tool user. Generalization beyond ecologically applicable skill is therefore limited. If capuchin nut-cracking skill is influenced by nut resistance, future research could investigate whether capuchins display improved skill with softer target nuts as capuchins are known to adjust percussive force in accordance with nut properties ^52,53^.. As this is outside the scope of this study we focus our comparative interpretations of skill exclusively on macaques and chimpanzees, as target nuts in both home ranges were of similar structural resistance.

Across all tested variables - foraging duration, cracking efficiency, success, and precision - long-tailed macaques are the more proficient nut crackers compared to chimpanzees. Macaques consistently take the least amount of time to forage and apply the fewest hits to successfully crack open the nuts, while rarely losing the nut through mishandling of the tool. Overall, macaques are the most successful forager, feeding on almost all nuts they attempt to crack (∼95%). Chimpanzees are less proficient when compared to macaques. Chimpanzees take more time to consume a nut and require more hits per nut than macaques. This shows that the strength of a primate species is not an important factor during nut cracking. Chimpanzees have previously been shown to use accumulated hits onto the nut to incrementally approach the breakage point of the nut ^34^.

Behavioural characteristics during nut cracking vary only slightly between macaques and chimpanzees. Both species crack nuts in a sitting position using one hand to guide the hammer. Macaques consistently engage in shielding and they generally strike the nut from a higher maximum height relative to their body size. Nonetheless, this variation in techniques might be responsible for the enhanced efficiency of macaques. We suggest that shielding plays a central role for precision and as a result for nut-cracking skill of macaques.

Furthermore, the relative skill of a macaques and chimpanzees, as described above, had an effect on the degree of percussive damage inflicted upon their tool materials. Even though macaques processed more nuts with their tools compared to chimpanzees, macaque hammerstones and anvil showed the lowest relative degree of macro percussive damage compared to chimpanzees. As raw material was controlled for, it can be suggested that variation in skill is related to surface modification on both stone hammers and anvils, with lower skill levels resulting to an increase in damage.

As the nut species used in this study are considerably softer than both the stone hammers and anvils it can be assumed that the observed damage is produced through accidental stone-on-stone contact between hammer and anvil ^21,23^. The degree of damage is likely related to imprecise striking where the hammerstone accidently strikes the anvil. Neither species showed a notable number of complete mishits, the stone-on-stone contact must therefore follow from strikes that hit both nut and anvil, as shown elsewhere ^21^.

The more often a tool is used, the higher the likelihood for damage to occur. Furthermore, the adjustment of tools during use additionally can affect the distribution of damage patterns across the stone. Due to circumstances in the field, each primate species did not crack the same number of nuts, resulting in both hammers and anvils being used disproportionately between species. Capuchins cracked about 1000 nuts per anvil, whilst it is estimated that macaque cracked between 600 and 700 nuts, and chimpanzee cracked around 450 nuts per anvil. Taking this variation into account, however, further highlights the effect of inter-species skill, especially between macaques and chimpanzees. Although macaques cracked a greater number of nuts compared to chimpanzees, they produced fewer damage on both hammers and anvils. Concluding on the comparison of capuchin and chimpanzee tools is more difficult, since it is unclear if chimpanzee hammers had shown similar amounts of use wear after additional use.

In addition to species-specific differences in overall amounts of use wear, hammerstones exhibit different damage distributions. Little and centralized damage on macaque hammerstones indicates a high level of homogeneity regarding how macaques handle the tools. They handled hammerstones in a way that would regularly affect the same areas of the tool surface, which suggests a high degree of control over hammer movement. In contrast, high amounts of use wear and a high-density distribution on capuchin hammerstones indicate little control over hammer motion, as well as high variability in hammer handling and high amounts of rotation of the tool during use. Use wear patterns on chimpanzee hammerstones suggest intermediate levels of homogeneity regarding hammer handling. As chimpanzees were never observed to adjust their grip on the hammer stone, increased number of use wear areas and of overall damage as compared to macaques can be ascribed to lower levels of efficiency and precision. Nut-cracking of each species thus correlates with a specific damage pattern that can be explained through differences in nut-cracking skill. Chimpanzees and capuchin monkeys are known to have cracked nuts for thousands of years ^54–56^. Until now, there is no evidence of long-term nut cracking behaviour in macaques. Due to aquatic environments at tool sites within tidal zones archaeological investigations have not yet been able to identify older tools ^57^. However, behavioural observations have shown that macaques display percussive stone tools use to forage for shellfish for at least a century ^58^. Macaque tool behaviours, in particular oyster cracking, require precise hammering in order to crack open the shell without damaging it. Since oysters are attached to substrate their position cannot be actively manipulated which requires macaques to work from different angles, sometimes overhead ^59^. Oyster cracking thus requires controlled and precise percussive movements with regards to where the oyster is hit and with how much force the strike is executed with. This precise form of tool-use has not been documented in any other non-human primate ^59^. Highly precise tool techniques might explain the levels of technical skill during nut cracking in long-tailed macaques.

An additional factor which may affect the material record of inter-species percussive behaviour is group composition. If greater numbers of individuals use the same hammer and anvil, the potential for a wider diversity of damage patterns to develop is higher, as each individual may manipulate the tools in different ways or exhibit varying levels of skill. In our study, chimpanzee and macaque tool-users comprised relatively low numbers (chimpanzee n=7 and macaque n=5) of individuals, while the greater number of capuchin monkeys (n=18) displayed a high diversity of tool-users, including juvenile individuals. However, juvenile participation in the nut-cracking experiment was marginal and most likely did not influence the overall damage pattern seen on capuchin tools, differing patterns of use-wear produced by individuals cannot be excluded, nor directly disentangled from the aggregate end use wear pattern on both hammers and anvils. A similar situation can, however, be suggested for natural hammerstones and anvils in the environment of tools users. Surface modifications on these artefacts are potentially a result of multiple different tool-users over varying periods of time ^60,61^.

The macaque and chimpanzee anvils and hammerstones did not exhibit extensive surface modifications which would make them difficult to identify as tools when found in the landscape. This follows from highly efficient and precise nut-cracking, and in the case of chimpanzees likely also from using the tools for a limited amount of time. Capuchin artefacts on the other hand display obvious battered areas, which are indicative of percussive tool use. The increased nut resistance of capuchins and the resulting imprecise striking with great force should result in clearly visible macro percussive damage. This study contributes to the referential framework of pounding tools that can inform the archaeological record as examples of percussive use wear of a foraging behavior. In conclusion, our findings suggest that the observed inter-species skill variation seemingly effects the degree at which percussive damage accrues on both hammerstones and anvils. Higher skill levels are suggested by an overall reduction of percussive damage. Our study suggests that the archaeological signature of highly skilled nut-crackers may be considerably less identifiable in the archaeological record of primates.

The use of percussive technology by Plio-Pleistocene hominins to access bone marrow is well documented through the identification of characteristic broken bones and bone fragments at Early Stone Age archaeological sites, often found in association with hammerstones ^18,62^. While it has been hypothesized that hominins engaged in a wider range of percussive behaviors, such as nut cracking, there is little direct evidence to support this. Archaeological examples of percussive anvils have been interpreted, through comparison with experimental data, as components in bipolar knapping ^7,63,64^.

This study has shown that more highly skilled nut crackers are less likely to leave a visible percussive signature on their stone tools. This suggests that Oldowan hominins possessed a high degree of manual dexterity, given the skill exhibited in many Oldowan assemblages ^65,66^. The same level of tool manipulation skill was likely employed during percussive activities. Consequently, the results of this study suggest that artefacts (anvils and hammers) associated with percussive behaviours, such as nut cracking, are likely underrepresented in the early stone age archaeological record because they are difficult to distinguish from natural, unmodified stones.

In cases where nut cracking leaves little in the form of a durable material signature, new methodologies must be developed that entail a high enough resolution to identify ephemeral percussive damage ^67^ while also differentiating this damage from taphonomic alterations on the stone surfaces^24^. However, since the degree of percussive damage seems related to the extent to which these stone tools are used, there is potential for such behaviours to be preserved as a material signature at sites that have seen repeated occupations over a prolonged period and reuse of the same, potentially larger artefacts, such as anvils, for the same behaviours.

## 2. Material and Methods

### 2.1 Material

Each species (chimpanzees, macaques and capuchin monkeys) was presented with a tool set consisting of an anvil and three hammerstones. Sizes and characteristics, such as material type, shape, weight and surface condition of anvil and hammer stones were similar for all three species. This ensured that damage pattern and behaviours would not be affected by differing tool properties. Weights were chosen to represent a range within the natural tool selection size of all species, comprising relatively small chimpanzee tools and roughly small to medium sized macaque and capuchin tools ^48,49^.

Sizes and characteristics, such as material, form, weight and surface condition of anvil and hammerstones were selected to be similar for all three set ups. Hammerstones provided were round igneous rocks (common in the Oldowan). Capuchin and macaque hammerstones were phonolite cobbles sourced from the Olduvai Gorge, Tanzania and chimpanzee hammers were basalt cobbles from Koobi Fora, Kenya. Raw material properties are roughly identical with materials sharing similar internal structures. Olduvai Gorge and Koobi Fora both comprise Oldowan sites, and phonolite and basalt are abundant in each location and part of the Oldowan record. Neither of these raw materials occur naturally in the respective primate habitats, ensuring that neither species was familiar with the material prior to exposure (see Supplementary Materials and Supplementary Table 1 for details on provided raw materials).

All primate species were provided with fresh palm nuts native to their habitats. Nuts presented to chimpanzees and macaques were oil palm nuts of the species *Elaeis guineensis*. The same nut species was initially provided to the capuchins. After initial tasting of the nut the capuchins however refused to eat this species of palm nuts and were therefore provided with palm nuts of the species *Syagrus romanzoffiana*, which are native to their habitat. Palm nuts are relatively low-resistance nuts ^26,68^ and all primate species in our study have previously been observed to forage on their local oil palm species using hammerstones and anvils. Between 30 to 50 nuts at a time were placed next to the anvil when the setup was prepared. The structural resistance of both nut species presented to each primate group was tested experimentally by dropping a 500g weight through a 50 cm tube nut, counting the number of drops required to successfully crack it open. We then compared the data across locations.

### 2.2 Methods

#### 2.2.1 Behavioral Analysis

Data collection took place in Bossou, Guinea for chimpanzees (*Pan troglodytes verus*), in the Tietê Ecological Park São Paulo, Brazil for capuchins (*Sapajus libidinosus*), and in the Ao Phang Nga National Park, Thailand for macaques (*Macaca fascicularis*). Bossou chimpanzees live in a habitat of 5 to 7 km^2^ around the village Bossou (for a detailed description of the habitat see ^69^). The nut-cracking setup was placed in the “outdoor laboratory” of the field site that lies within the core range of the population. Capuchins who participated in this study live in a semi-wild environment of 14 km^2^ within Tietê Ecological Park (for information on vegetation and provisioning see ^70^). The long-tailed macaques in this study live in the southern end of Boi Island, Phang Nga National Park. This group is not habituated to human observers. Oil palm was introduced into their habitat by humans in the early 2000s ^30^. (For detailed information on participating individuals see Supplementary Table 2).

Each species was presented with the same nut-cracking setup with the final aim of cracking a total of 1000 nuts each. Chimpanzee data was collected by a handheld video camera and camera traps over the course of nine days in November and December 2018, resulting in 165 minutes of video material. Both data collection types led to videos from which tool selection and use could not be fully identified, due to a blocked view or too great distance from the camera. While only 140 events could be coded, based on how many nuts chimpanzees were provided, they are estimated to have cracked about 450 nuts with the provided hammerstones and anvils. Capuchin video data was recorded during four days in April 2017 during direct observation of the monkeys, leading to 5h of video data. Video data consistently, and with few exceptions, covered the full timespan and details of the event, resulting in 945 nut cracking events. Macaque video data was recorded by camera traps in January and February 2017 over the course of seven days. Macaques were provisioned with 1000 nuts at the anvil, from which 485 individual nut cracking events could be identified on video. To describe and compare the behaviour between species when nut cracking, we assessed “Posture” (sitting/standing), “Hammer Height” (relative to body size and posture), “Handedness” (right/left/both/switch), “Shielding” (hand as barrier next to nut), and “Cracking” (nut cracked but not necessarily eaten) (For a detailed description of all variables see Supplementary Material: Methods 2).

To compare nut cracking skill, we investigated four different components:

##### 1. Foraging Duration

We first compared nut-cracking durations (time needed to crack and consume a single nut) across species. The less time needed per nut would yield an overall greater foraging efficiency over the same amount of time spent nut-cracking. This will be used to indicate efficiency differences between species.

##### 2. Cracking Efficiency

Here we assessed the number of hits each species needs to crack open a nut. Due to the different nut-cracking techniques applied by each species, this measure does not directly translate to energetic cost. However, knowing the number of hits each primate requires to crack a nut helps to form a complete picture of the nut-cracking process. Taken together with the other behavioural measures, this contributes to assessing the species-specific efficiency.

##### 3. Success

We compared the successful cracking events to failed attempts for each species. Nut cracking events were counted as successful when the primate was able to eat the nut or at least parts of it.

##### 4. Striking Precision

Less precise hits are characterized by strikes that contact the nut at an angle that forces it off the anvil. Thus, to assess precision levels of each species, we compared the number of times that the nut was displaced off the anvil in relation to the total number of hits during one nut-cracking event. Whenever the hammerstone is in direct contact with the anvil, damage may occur on the tool materials. Use damage therefore can be used to assess nut cracking precision.

#### 2.2.2 Statistical analysis

We carried out separate statistical analyses for each component of technical skill (Foraging Duration, Cracking Efficiency, Success, and Precision). All analyses were conducted in R (V4.0.5, ^71^). Skill was analysed using functions lmer and glmer of the statistics package lme4 ^72^ to run linear mixed models (LMM) and generalized linear mixed models (GLMM; ^73^). All analyses include only subadult and adult individuals of both sexes for whom five or more observations were available for the respective model. Due to the small numbers of individuals per species in some cases we could not assess the influence of age and sex.

Response variables for each model were “Duration” (for Foraging Duration), “Hits” (for Cracking Efficiency), “Success”, and “Precision” (for Striking Precision). To investigate if species differ in the time it takes to crack and consume one nut (“Duration”), we used an LMM with Gaussian error structure and identity link function. We log-transformed the response prior to running the model to achieve an approximately symmetrical distribution and to avoid potentially influential cases. To investigate if species differ with regards to how many hits were needed to crack a single nut (“Hits”), we used a GLMM with Poisson error structure and log link function. To assess differences between species regarding how successful (“Success”) and how precise (“Precision) they crack nuts, we used GLMMs with binomial error structure and logit link function. For the “Success” model, the response was the binary variable “Success” (nut was consumed or not consumed), transformed to a numerical variable with 0 for “not successful” and 1 “successful”. The “Precision” model response was stored as a matrix of (Nut displaced from anvil, Hits per Nut – Nut displaced from anvil) to take total number of hits per event into account. Models for “Cracking Efficiency”, “Success”, and “Precision” were fitted using the optimizer “bobyqa”.

Each model comprised the response variable of interest, “Species”, as a fixed effect and controlled for hammer selection by including the fixed effect “Hammer Size” (three different sizes were presented to each species). We further included “Subject” (which is the individual primate), “Hammer ID”, and “Hammer Size” nested in “Subject” as random intercept effects. The “Precision” model additionally included observation level random effects (“Nut.ID”) to deal with a non-binary response in a model with binomial error structure. Control and random predictors were included to keep type I error rate at the nominal level of 5% ^74^.

For the Gaussian model on Foraging Duration, we visually checked normality and homogeneity of residuals. Both checks revealed no violation of these assumptions. We ruled out overdispersion for the models Cracking Efficiency and Precision. For all models, we visually inspected distributions of the “Best Linear Unbiased Predictors” ^73^. All models revealed sufficiently normally distributed random effects. We further assessed model stability for all models by comparing estimates of the original model to estimates of models obtained by dropping one level of each random effect grouping factor at a time ^75^. All models reveal stable model estimates, with relatively wide ranges for the Success and Precision model estimates. To assess the role of individual subjects in more detail, we further investigated influential cases of the random effect “Subject” with a Cook’s Distance plot of the statistical package influence.ME ^75^ for each model. Visual inspection revealed one to two potential influential cases per model. To assess the gravity of their influence, we compared model estimates of the original models to estimates of models obtained after excluding potential influential cases. This suggests no strong influence of single individuals for all models, but the Success model. The particularly low success rate (mean=0) of an adult female capuchin affects the significance level on differences between species. Excluding her from the Success model renders the difference between capuchins and chimpanzees no longer significant (p=0.0598). However, our analysis is focused on assessing the overall effect of the variable “Species”, interpretation of results is not based on pairwise comparisons between species. The overall effect of species remains relevant when excluding the respective individual. We kept complete datasets for all models. To rule out collinearity, we calculated the Variance Inflation Factor (VIF; ^76^) for a standard linear model with fixed effects only. See Supplementary Materials: Methods 3. for a detailed description of assumption checks.

Confidence intervals of model estimates were calculated using the function bootMer of the package lme4 by bootstrapping over the full model using 1,000 parametric bootstraps. To avoid cryptical multiple testing, an overall effect of “Species” for each model was assessed by a likelihood ratio test, comparing the full model to a reduced model lacking the variable “Species” but with otherwise identical model structure as the full model ^77,78^. Effect sizes ^79^ were calculated using the function r.squaredGLMM of the statistical package MuMin ^80^.

Sample sizes varied between models, with 824 observations available for the Foraging Duration Model, 870 observation for the Cracking efficiency model, 1,508 observations for the Success model and 1,459 observations for the Precision model. (For more details see Supplementary material “Methods” section).

#### 2.2.3 2D and 3D GIS data collection of percussive damage on stone tools

To analyse use wear patterns on hammerstones, we applied 2D and 3D GIS procedures ^23,25^. 3D models of all nine hammerstones were generated using 216 photographs per hammerstone processed in the photogrammetry software Agisoft Metashape (Version 1.6.5). This produced a point density cloud of 25,000 points per cm^2^. Photographs were taken with a Nikon D850, a 50-megapixel sensor and by using a Foldio 360 automated turntable with a 10-degree interval. The process of building the models was semi-automated using several custom python scripts, for model meshing, model texturing and to build sparse and dense point clouds ^25^.

Assessment of surface damage on all hammerstones was done in two phases, the first, on the uncleaned hammerstones, where less developed surface modification was the most visible. A second assessment was conducted following a thorough washing of the hammerstones to ensure that all surface modification was identified. Photorealistic 3D models of the cleaned hammerstones were processed in the open access 3D software Blender (V2.90.1, ^81^). All traces of percussive damage were extracted by hand for each hammerstone in Blender. To ensure accuracy, contiguous damaged areas were first visually inspected on the original hammerstones and then identified on and extracted from the 3D model. (For a detailed description of the Blender workflow see ^25^). Modified and unmodified surface areas were exported for further analysis. We calculated surface area, perimeter and maximum dimension for each use wear segment, the following were analysed via the 3D modelling library PyVista ^82^ in Python. Variables comprised percentage of total use wear area (PA), the largest single use wear area (LUW), the density of use wear on the hammerstone (D), and the density of use wear in relation to the total surface area (ED).

To quantify varying degrees of anvil damage between species, the active surface of each anvil was subjected to a 2D GIS analysis (applying methods outlined in ^23^). Images of each active surface were georeferenced using a local coordinate system using QGIS ^83^. The total surface area of the active plane of each anvil was calculated. Each contiguous area of percussive damage was outlined and converted to a separate shapefile. These were used to calculate the degree of percussive damage in relation to the total surface area of the active plane and total perimeter of all use wear areas.

## Supporting information

Luncz_Slania2024_Supplementary_information

## Acknowledgements

We thank the Centro de Resgate de Animais Silvestre (CRAS) from the Tietê Ecological Park (PET) in São Paulo, Brazil, the National Research Council of Thailand and the Institut de Recherche Environmentale de Bossou (IREB) of the Republic of Guinea for enabling our research (31/MESRS/DGERSIT/2018). We thank the Department of National Parks, Wildlife and Plant Conservation for permission to conduct research in Ao Phang Nga National Park. We thank the park head Mr. Roengnarong Nimnuan for supporting our work and granting access to the park. Research in Guinea was granted by the Direction Générale de la Recherche Scientifique et de l’Innovation Technologique (31/MESRS/DGERSIT/2018). Raw material collection at Olduvai was authorized by the Commission for Science and Technology (COSTECH), the Ngorongoro Conservation Area Authority, and the Department of Antiquities, Tanzania under the following COSTECH permits (BICAEHFID, ERC-AdvG 832980). Material collection in Koobi Fora was authorized by the National Museums of Kenya (NACOSTI/P/18/28541/23258).

This research was funded by the German Primate Center in Göttingen and the Max Planck Society. During analysis T.P was funded by a British Academy Fellowship (Project Number: 542133). A.A was supported by the Spanish Ministry of Science, Innovation, and Universities through the Ramón y Cajal Grant (RYC2022-037569-I) funded by MICIU/AEI/10.13039/501100011033 and by ESF. K.A.W was supported by the Fundação para a Ciência e Tecnologia (SFRH/BD/115085/2016), the National Geographic Society (EC-399R-18), and The Leverhulme Trust (ECF-2022-322). T.F. was funded by Fundação de Amparo a Pesquisa do Estado de São Paulo (#2018/01292-9).

## Data availability statement

All code generated is provided in the Supplementary Materials. The original data sets are available upon request.

## Author contributions

Conceptualization: LVL, TP Data curation: LVL, NES, TP

Formal analysis: NES, LVL, AA, TP Funding acquisition: LVL

Methodology: LVL, NES, KAW, TF, TP Project administration: LVL

Resources: LVL, SC, TF, IDLT, SM, TP

Supervision: LVL Visualization: NES, TP

Writing - original draft: NES, LVL

Writing - review & editing: LVL, NES, KAW, SC, TF, SM, TP

