## Supplementary material for "Tool skill impacts the archaeological evidence across technological primates": Luncz_Slania2024_Supplementary_information

[ꭗ]joint 1[st] author

**Supplementary Material:**

### Materials

1. Stone material provided in experimental setup:

Hammers systematically differed in weight and size within an experimental setup, making for a small (257.6-367.6g), medium (304.5-462.6 g) and large (580.6-661.5 g) hammerstones in each set up (Supplementary Table 1). Weights were chosen as to represent a range that falls within the preferred tool size of all species, comprising relatively small chimpanzee tools and roughly small to medium sized macaque and capuchin tools. All anvils were quartzite tabular blocks. Capuchin and macaque anvils were imported from Olduvai Gorge and chimpanzee anvil stones from Koobi Fora.

**Supplementary Table 1:**
Weights of each hammerstone before use.

| **Species** | **Size** | **Weight in gr** |
|---|---|---|
| Chimpanzee | small | 339.6 |
| Chimpanzee | medium | 462.6 |
| Chimpanzee | large | 629.9 |
| Capuchin | small | 257.6 |
| Capuchin | medium | 304.5 |
| Capuchin | large | 580.6 |
| Macaque | small | 367.6 |
| Macaque | medium | 441.1 |
| Macaque | large | 661.5 |

### Methods:

1. Observational data collection

**Supplementary Table 2:**
Participating individuals in the comparative nut cracking experiments per Species, Sex, and Age. Juvenile individuals were excluded from behavioral analysis.

| | **Chimpanzees** | **Capuchins** | **Macaques** |
|---|---|---|---|

| | M | F | M | F | M | F |
|---|---|---|---|---|---|---|
| Adult | 2 | 3 | 3 | 3 | 3 | 1 |
| Subadult | - | 1 | 5 | - | 1 | - |
| Juvenile | - | - | 6 | 1 | - | - |
| **Total** | 6 | | 18 | | 5 | |

2. Behavioural variables coded

For each event, we identified **Subject** and **Species**, **Sex**, and **Age Group** of the individual. The cracking or attempted cracking of one nut was coded as one event and all of the variables were assessed for each nut-cracking event, i.e. one nut, whenever feasible. When camera angle, video quality or video timeframe would not allow to extract all variables from one event, all visible variables were coded, and not visible variables were left out for this event.

- **Nut Duration**: The time it took an individual to crack and (if successful) eat one nut. The event starts with the individual securing the nut. It ends when the individual is done eating the nut, when the individual selects a new nut, when the individual gets up to leave or when the individual stops cracking.
- **Hammer Size**: Size (small, medium, large) of the hammer that individual choses.
- **Hammer ID**: ID of hammer used.
- **Hits per Nut**: Total number of hits of the event, including hits on the nut and mishits.
- **Anvil Hits**: Nr of times the individual hits the anvil.
- **Mishits**: Nr of times the individual misses the nut.
- **Nut Flying Off**: Number of times the nut jumps off when hit with the hammer during one nut-cracking event. This includes the nut rolling away.
- **Success**: records whether the individual eats the nut or not.
- **Cracking**: If the individual cracked the nut open or not. If the event ends with the individual leaving, a disturbance, the nut jumping away etc. and the individual did not crack the nut yet, the nut is considered not cracked. Only if we do not see the end of the event, e.g. when a video stops or the view is blocked, it is unknown.
- **Handedness**: Hand the individual uses to strike the hit (right, left, both, switch).
- **Shielding**: The individual is shielding the nut on the anvil using one hand as a barrier next to the nut so the nut cannot fly off.
- **Posture**: differentiates whether the individual is sitting or standing upright or standing bent when raising the stone.
- **Hammer Height**: Records how high the individual raises the hammer in relation to the body (Belly, Chest, Shoulder, Forehead, Overhead). The highest point of all hits is being coded.
- **Adjustments:** Number of times an individual adjusts their grip on the hammer stone during one cracking event.

3. Statistical Analysis

Assumption checks for statistical models

<u>Foraging Duration model:</u>

The sample for this model consisted of 824 observations of 17 individuals: 62 observations of 6 chimpanzees, 391 observations of 7 capuchins, and 371 observations of 4 macaques.

We visually inspected a histogram and qqplot to assess normality of residuals (Field, 2005) and plotted fitted values against residuals to check homogeneity of residuals (Quinn and Keough, 2002;). This revealed normally distributed and homogeneous residuals. We further visually inspected the distribution of the “Best Linear Unbiased Predictors” (BLUPs; Harrison et al. 2018). We judged the distribution of BLUPs to be adequate. We assessed model stability by comparing estimates of the original model to estimates of models obtained by dropping one level of the random effect Subject at a time (Nieuwenhuis, Grotenhuis, & Pelzer, 2012). This revealed stable model estimates (see Supplementary Table 3). We further investigated the role of individual subjects by visually inspecting a Cook's Distance plot (Supplementary Figure 3) using the R package "influence.ME" (Nieuwenhuis, Grotenhuis, & Pelzer, 2012). This suggests that the chimpanzee "Yo" could be influential. Yo is a particularly old (57 years) female chimpanzee. She needed more time to crack a single nut (mean=75.7s), as compared to the groups average excluding here (mean=27.61s). Excluding Yo from the dataset affects model estimates slightly. Importantly, significance levels remain unaffected by her exclusion. We kept the full dataset. To rule out collinearity we assessed the variance inflation factor with the R package "car" (Field, 2005). This revealed no issues with collinearity ($GVIF^{(1/(2*df))}$=1.13).

**Supplementary Table 3:**
Model estimates, std. errors, degrees of freedom (df), t-values, and p-values of (A) the original model and (B) the model after excluding influential cases. Model to assess foraging duration.

| **A** | Estimate | Std. Error | df | t | P |
|---|---|---|---|---|---|
| (Intercept) | 2.870 | 0.153 | 11.771 | 18.736 | 0.000 |
| SpeciesChimpanzee | 0.348 | 0.242 | 12.644 | 1.441 | 0.174 |
| SpeciesMacaque | -0.986 | 0.253 | 11.231 | -3.896 | 0.002 |
| HammerMedium | -0.012 | 0.107 | 6.702 | -0.113 | 0.914 |
| HammerSmall | 0.120 | 0.122 | 6.353 | 0.981 | 0.362 |
| **B** | Estimate | Std. Error | df | t | P |
| (Intercept) | 2.861 | 0.129 | 11.603 | 22.212 | 0.000 |
| SpeciesChimpanzee | 0.146 | 0.218 | 12.979 | 0.668 | 0.516 |
| SpeciesMacaque | -0.989 | 0.213 | 11.443 | -4.632 | 0.001 |
| HammerMedium | 0.010 | 0.097 | 6.323 | 0.101 | 0.923 |
| HammerSmall | 0.123 | 0.111 | 6.051 | 1.107 | 0.310 |

Cracking Efficiency model:

This model made use of a sample of 870 observations of 16 individuals: 65 observations of 5 chimpanzees, 419 observations of 7 capuchins, and 395 observations of 4 macaques.

We visually inspected the distribution of BLUPs which revealed normally distribution of random effects. Model stability was checked by comparing estimates of the original model to estimates of models obtained by dropping one level of each random effect at a time. All model estimates were stable. We further investigated influential cases by visually inspecting a Cook's Distance plot (see Supplementary Figure 5). This indicates that the chimpanzee Yo, again, and to a lesser extent the capuchin Frida could be influential. Yo needed a greater average number of hits (mean=10.88) to crack a single nut compared to the other chimpanzees (mean=6.58). The capuchin Frida is an adult female who needed a greater mean number of hits to crack a nut (mean=3.17), as compared to average number of hits of capuchins excluding Frida (mean=1.73). Excluding both individuals from the dataset notably affects model estimates, std. errors and p-values, without influencing direction of effects or significance levels (see Supplementary Table 6). We kept the complete dataset.

We assessed potential collinearity by calculating the variance inflation factor based on a model with fixed effects only, which revealed no collinearity issue ($GVIF^{(1/(2*df))}$=1.12). With an dispersion parameter of 0.59 ($X^2$= 511.55, df=862, p=1) the model is underdispersed, rendering it conservative.

**Supplementary Table 4:**
Original model estimates in comparison with minimum and maximum estimate values from dropping random effects one at a time. Model to assess cracking efficiency.

| | orig | min | max |
|---|---|---|---|
| (Intercept) | 0.578 | 0.518 | 0.900 |
| SpeciesChimpanzee | 1.208 | 0.935 | 1.315 |
| SpeciesMacaque | -0.589 | -0.798 | -0.454 |
| HammerMedium | 0.281 | 0.099 | 0.384 |
| HammerSmall | 0.319 | 0.156 | 0.509 |
| Hammer.in.Subj@(Intercept)@NA | 0.129 | 0.000 | 0.151 |
| Subject@(Intercept)@NA | 0.217 | 0.136 | 0.308 |
| Hammer.ID@(Intercept)@NA | 0.000 | 0.000 | 0.111 |

**Supplementary Table 5:**
Model estimates, std. errors, z-values, and p-values of (A) the original model and (B) the model after excluding influential cases. Model to assess cracking efficiency.

| **A** | Estimate | Std. Error | z value | Pr(>\|z\|) |
|---|---|---|---|---|
| (Intercept) | 0.578 | 0.106 | 5.446 | 0.000 |
| SpeciesChimpanzee | 1.208 | 0.160 | 7.535 | 0.000 |
| SpeciesMacaque | -0.589 | 0.182 | -3.227 | 0.001 |
| HammerMedium | 0.281 | 0.094 | 2.999 | 0.003 |
| HammerSmall | 0.319 | 0.120 | 2.653 | 0.008 |
| **B** | Estimate | Std. Error | z value | Pr(>\|z\|) |
| (Intercept) | 0.505 | 0.077 | 6.563 | 0.000 |
| SpeciesChimpanzee | 1.144 | 0.120 | 9.546 | 0.000 |
| SpeciesMacaque | -0.464 | 0.128 | -3.619 | 0.000 |
| HammerMedium | 0.317 | 0.090 | 3.513 | 0.000 |
| HammerSmall | 0.229 | 0.118 | 1.935 | 0.053 |

<u>Success model:</u>

This model was based on a sample of 1,508 observations of 20 individuals: 125 observations of 5 chimpanzees, 938 observations of 10 capuchins, and 445 observations of 5 macaques.

We assessed the distribution of BLUPs visually which revealed normally distributed random effects, with the exception of the random grouping factor Hammer.ID. We investigated model stability by comparing estimates of the original model to estimates of models obtained by dropping one random effect at a time. This revealed mostly stable model estimates with some uncertainty regarding the Intercept and the estimate for small hammers (see Supplementary Table 7). To assess influential cases in more detail, we visually inspected a Cook's Distance plot for the random effect "Subject". This suggests that the capuchin Cisca and to a lesser extent the capuchin Vodka are influential. The adult female Cisca did not successfully consume a single nut, while the adult male Vodka shows a higher success rate (0.53) compared to the average success of all capuchins excluding Vodka and Cisca (0.46). Excluding both subjects from the datasets, greatly influences model estimates with the estimated term for the intercept changing direction and the estimate for species chimpanzee no longer being significant (see Supplementary Table 8), meaning that the difference in success between chimpanzees and capuchins no longer reaches significance. The extraordinary low success rate of the adult female capuchin Cisca therefore greatly impacts overall capuchin success rate in comparison to the other species. While a pairwise comparison of the variable “Species” is thus affected by excluding influential cases, the overall species effect remains significant (Chisq=18.83397, Df=2, p=0.0001). We kept all individuals in the dataset, but the influence of Cisca should be considered when evaluating results. To rule our collinearity, we calculated the VIF based on a model only including fixed effects. This revealed no issue with collinearity ($GVIF^{(1/(2*df))}$=1.08)

**Supplementary Table 6:**

Original model estimates in comparison with minimum and maximum estimate values from dropping random effects one at a time. Model to assess success.

| | orig | min | max |
|---|---|---|---|
| (Intercept) | -0.334 | -0.511 | 0.748 |
| SpeciesChimpanzee | 1.509 | 0.259 | 2.263 |
| SpeciesMacaque | 3.558 | 3.070 | 4.324 |
| HammerMedium | -0.416 | -1.236 | -0.065 |
| HammerSmall | -0.485 | -1.347 | 0.038 |
| Hammer.in.Subj@(Intercept)@NA | 0.417 | 0.000 | 0.656 |
| Subject@(Intercept)@NA | 0.783 | 0.000 | 0.938 |
| Hammer.ID@(Intercept)@NA | 0.179 | 0.000 | 0.497 |

**Supplementary Table 7:**

Model estimates, std. errors, z-values, and p-values of (A) the original model and (B) the model after excluding influential cases. Model to assess cracking efficiency.

| **A** | Estimate | Std. Error | z value | Pr(>\|z\|) |
|---|---|---|---|---|
| (Intercept) | -0.334 | 0.408 | -0.818 | 0.413 |
| SpeciesChimpanzee | 1.509 | 0.662 | 2.279 | 0.023 |
| SpeciesMacaque | 3.558 | 0.636 | 5.591 | 0.000 |
| HammerMedium | -0.416 | 0.352 | -1.181 | 0.237 |
| HammerSmall | -0.485 | 0.441 | -1.100 | 0.271 |
| **B** | Estimate | Std. Error | z value | Pr(>\|z\|) |
| (Intercept) | 0.017 | 0.313 | 0.054 | 0.957 |
| SpeciesChimpanzee | 0.858 | 0.456 | 1.882 | 0.060 |
| SpeciesMacaque | 3.464 | 0.479 | 7.227 | 0.000 |
| HammerMedium | -0.380 | 0.338 | -1.125 | 0.261 |
| HammerSmall | -1.007 | 0.446 | -2.256 | 0.024 |

<u>Striking Precision model:</u>

The sample for this model encompassed 1,459 observations of 17 individuals: 115 observations of 4 chimpanzees, 938 observations of 10 capuchins, and 406 observations of 3 macaques.

We assessed the distribution of BLUPs visually which revealed normally distributed random effects, with the exception of the random grouping factor Hammer.ID. Model stability was assessed by comparing model estimates of the original model to estimates of models obtained by dropping one random effect at a time. This revealed stable model estimates with the exception of estimated terms regarding the variable Hammer (see Supplementary Table 9). To assess influence of individual subjects, we visually inspected a Cook's Distance plot. This revealed potential influence of the adult male macaque Bob. Bob's rate of the nut rolling of the working surface to number of hits is slightly lower (0.1) as the group's average rate excluding him from the dataset (0.11). While model estimates change slightly, significance levels are not affected by excluding Bob from the dataset (see Supplementary Table 10). We kept a complete dataset for the model. To ensure there was no issue of collinearity we calculated the VIF for a standard linear model. This revealed VIFs of $GVIF^{(1/(2*df))}$=1.07 for Species and Hammer and hence no collinearity between variables. To rule out overdispersion in relation to the continuous response and binomial error structure, we calculated the dispersion parameter. This revealed a dispersion parameter of 0.88 (Chisq=1278.85, df=1450) and showed slight underdispersion.

**Supplementary Table 8:**
Original model estimates in comparison with minimum and maximum estimate values from dropping random effects one at a time. Model to assess precision.

| | orig | min | max |
|---|---|---|---|
| (Intercept) | -0.725 | -1.511 | -0.517 |
| SpeciesChimpanzee | -1.120 | -1.631 | -0.572 |
| SpeciesMacaque | -2.043 | -2.627 | -1.668 |
| HammerMedium | 0.041 | -0.257 | 0.669 |
| HammerSmall | 0.242 | -0.360 | 1.019 |
| Obs.ID@(Intercept)@NA | 0.477 | 0.311 | 0.702 |
| Hammer.in.Subj@(Intercept)@NA | 0.000 | 0.000 | 0.297 |
| Subject@(Intercept)@NA | 0.373 | 0.000 | 0.433 |
| Hammer.ID@(Intercept)@NA | 0.256 | 0.000 | 0.311 |

**Supplementary Table 9:**
Model estimates, std. errors, z-values, and p-values of (A) the original model and (B) the model after excluding influential cases. Model to assess precision.

| **A** | Estimate | Std. Error | z value | Pr(>\|z\|) |
|---|---|---|---|---|
| (Intercept) | -0.725 | 0.270 | -2.689 | 0.007 |
| SpeciesChimpanzee | -1.120 | 0.374 | -2.992 | 0.003 |
| SpeciesMacaque | -2.043 | 0.401 | -5.101 | 0.000 |
| HammerMedium | 0.041 | 0.277 | 0.147 | 0.883 |
| HammerSmall | 0.242 | 0.338 | 0.716 | 0.474 |
| **B** | Estimate | Std. Error | z value | Pr(>\|z\|) |
| (Intercept) | -0.544 | 0.164 | -3.323 | 0.001 |
| SpeciesChimpanzee | -1.308 | 0.298 | -4.389 | 0.000 |
| SpeciesMacaque | -1.798 | 0.452 | -3.974 | 0.000 |
| HammerMedium | 0.039 | 0.150 | 0.257 | 0.797 |
| HammerSmall | -0.224 | 0.232 | -0.967 | 0.334 |

**Results Section:**

Description of behavioural techniques:

**Supplementary Table 10:** Behavioural details of nut cracking of chimpanzees, capuchins, and macaques. “Nr of Ind” states the number of individuals who showed the indicated behavioural variation at least once. “Nr of Events” refers to the absolute number of nut-cracking events. If an individual showed two or more behavioural variations, that it is included in all respective rows. Behavioural details are “Posture” during nut cracking, height of hammer relative to an individual’s body and posture (“Hammer Height”); hands used for cracking (“Handedness”), where “Switch” refers to a change from at least one variation to another within one cracking event, using a hand as a barrier next to the nut (“Shielding”); hammer stone grip adjustments (“Adjustments”); and the relative amount of nuts eaten after successful cracking (“Consumption of Nuts”). Percentage values are calculated based on all events for which the relevant measures were available and rounded to two decimal places. “Consumptions of nuts” is based on data of all individuals included in the respective model. Note that the percentage is based on those events for which the respective information was available.

| | | **Chimpanzee** | | **Capuchin** | | **Macaque** | |
|---|---|---|---|---|---|---|---|
| Total Nr of Ind | | | 6 | | 11 | | 5 |
| **Detail** | Variation | % of Events | Nr of Ind | % of Events | Nr of Ind | % of Events | Nr of Ind |
| Posture | Sitting | 98.39 | 6 | 7.55 | 5 | 99.39 | 5 |
| | Bipedal | 1.61 | 3 | 92.45 | 11 | 0.61 | 2 |
| Hammer Height | Overhead | 0 | 0 | 19.56 | 4 | 0.2 | 1 |
| | Forehead | 0.44 | 1 | 47.76 | 9 | 2.85 | 3 |
| | Shoulder | 11.8 | 2 | 25.90 | 11 | 43.38 | 5 |
| | Chest | 71.43 | 6 | 6.56 | 7 | 44.81 | 5 |
| | Abdominal | 16.33 | 5 | 0.22 | 1 | 0.2 | 1 |
| Handedness | Left | 90.27 | 4 | 0.21 | 2 | 35.63 | 3 |
| | Right | 9.73 | 2 | 5.45 | 1 | 64 | 2 |
| | Switch | 0 | 0 | 0.31 | 1 | 0.2 | 1 |
| | Both | 0 | 0 | 94.03 | 11 | 0.2 | 1 |
| Shielding | Yes | 5.84 | 3 | 6.08 | 2 | 62.41 | 5 |
| | No | 94.16 | 6 | 93.92 | 11 | 37.59 | 4 |
| Adjustments | Yes | 0 | 0 | 27.3 | 10 | 3.90 | 3 |
| | No | 100 | 6 | 72.7 | 10 | 96.1 | 5 |
| **Detail** | Variation | % of Events | Nr of Events | % of Events | Nr of Events | % of Events | Nr of Events |
| Consumption of Nuts | Yes | 60.60 | 77 | 45.00 | 442 | 94.38 | 420 |
| (Success) | No | 38.40 | 48 | 55.00 | 516 | 5.62 | 25 |

Foraging Duration model:

**Supplementary Table 11:**
Results of the log-duration model: Estimates with standard error, 95% confidence intervals, significance tests results.

| term | estimate | SE | lower CI | upper CI | t | df | P |
|---|---|---|---|---|---|---|---|
| Intercept | 2.870 | 0.153 | 2.567 | 3.152 | 18.736 | 11.772 | 0.000 |
| Species Chimpanzee | 0.348 | 0.242 | -0.121 | 0.834 | 1.441 | 12.645 | 0.174 |
| Species Macaque | -0.986 | 0.253 | -1.490 | -0.468 | -3.896 | 11.232 | 0.002 |
| Hammer Medium | 0.120 | 0.122 | -0.222 | 0.193 | 0.981 | 6.353 | 0.362 |
| Hammer Small | -0.012 | 0.107 | -0.124 | 0.374 | -0.113 | 6.702 | 0.914 |

**Supplementary Table 12:**
Model summary output of random effects of the foraging duration model, showing SD of each random intercept effect contributing to the response.

| Groups | Name | Variance | Std.Dev. |
|---|---|---|---|
| Hammer.in.Subj | (Intercept) | 0.03295 | 0.1815 |
| Subject | (Intercept) | 0.12669 | 0.3559 |
| Hammer.ID | (Intercept) | 0.00000 | 0.0000 |

<u>Cracking Efficiency model:</u>

**Supplementary Table 13:**
Results of the Cracking Efficiency model (Nr. of Hits): Estimates with standard error, 95% confidence intervals, and significance tests results.

| term | estimate | SE | lower CI | upper CI | z | P |
|---|---|---|---|---|---|---|
| Intercept | 0.578 | 0.106 | 0.353 | 0.799 | 5.446 | 0.000 |
| Species Chimpanzee | 1.208 | 0.160 | 0.882 | 1.529 | 7.535 | 0.000 |
| Species Macaque | -0.589 | 0.182 | -0.924 | -0.212 | -3.227 | 0.001 |
| Hammer Medium | 0.281 | 0.094 | 0.107 | 0.463 | 2.999 | 0.003 |
| Hammer Small | 0.319 | 0.120 | 0.073 | 0.547 | 2.653 | 0.008 |

**Supplementary Table 14:**
Model summary output of random effects of the cracking efficiency model, showing SD of each random intercept effect contributing to the response.

| Groups | Name | Variance | Std.Dev. |
|---|---|---|---|
| Hammer.in.Subj | (Intercept) | 0.0166729 | 0.1291237 |
| Subject | (Intercept) | 0.0468904 | 0.2165418 |
| Hammer.ID | (Intercept) | 0.0000000 | 0.0000002 |

Success model:

**Supplementary Table 15:**
Results of the Success model: Estimates with standard error, 95% confidence intervals, significance tests results.

| term | estimate | SE | lower CI | upper CI | z | P |
|---|---|---|---|---|---|---|
| Intercept | -0.334 | 0.408 | -1.078 | 0.395 | -0.818 | 0.413 |
| Species Chimpanzee | 1.509 | 0.662 | 0.404 | 2.733 | 2.278 | 0.023 |
| Species Macaque | 3.558 | 0.637 | 2.432 | 5.074 | 5.588 | 0.000 |
| Hammer Medium | -0.416 | 0.352 | -1.129 | 0.252 | -1.181 | 0.238 |
| Hammer Small | -0.485 | 0.441 | -1.409 | 0.341 | -1.100 | 0.271 |

**Supplementary Table 16:**
Model summary output of random effects of the Success model, showing SD of each random intercept effect.

| Groups | Name | Variance | Std.Dev |
|---|---|---|---|
| Hammer.in.Subj | (Intercept) | 0.17386 | 0.4170 |
| Subject | (Intercept) | 0.61349 | 0.7833 |
| Hammer.ID | (Intercept) | 0.03188 | 0.1785 |

Striking Precision model:

**Supplementary Table 17:**
Results of the Precision model (Nr. of Times Nut flies away in relation to number of hits): Estimates with standard error, 95% confidence intervals, significance tests results.

| term | estimate | SE | lower CI | upper CI | z | P |
|---|---|---|---|---|---|---|
| Intercept | -0.725 | 0.270 | -1.268 | -0.224 | -2.689 | 0.007 |
| Species Chimpanzee | -1.120 | 0.374 | -1.837 | -0.390 | -2.992 | 0.003 |
| Species Macaque | -2.043 | 0.401 | -2.849 | -1.257 | -5.101 | 0.000 |
| Hammer Medium | 0.041 | 0.277 | -0.506 | 0.636 | 0.147 | 0.883 |
| Hammer Small | 0.242 | 0.338 | -0.396 | 0.895 | 0.716 | 0.474 |

**Supplementary Table 18**:
Results of likelihood ratio tests of full-null model comparisons of each model (Foraging Duration, Cracking Efficiency, Success, and Precision).

| Effect: Species | $X^2$ | df | p |
|---|---|---|---|

| | | | |
|---|---|---|---|
| Foraging Duration | 16.85 | 2 | **0.0002** |
| Cracking Efficiency | 25.94 | 2 | **<0.0001** |
| Success | 17 | 2 | **0.0002** |
| Precision | 14.85 | 2 | **0.0006** |

**Supplementary Table 19:**

Model summary output of random effects of the Precision model, showing SD of each random intercept effect.

| Groups | Name | Variance | Std.Dev. |
|---|---|---|---|
| Obs.ID | (Intercept) | 0.22739 | 0.4769 |
| Hammer.in.Subj | (Intercept) | 0.00000 | 0.0000 |
| Subject | (Intercept) | 0.13901 | 0.3728 |
| Hammer.ID | (Intercept) | 0.06542 | 0.2558 |

### Use Wear

**Supplementary Figure 1:**
Use wear on chimpanzee hammerstones. Left to right: Plane A1, plane B1, and plan A2 of each stone. Small, medium and large hammer (top to bottom). Grey square: Scale of 10mm.

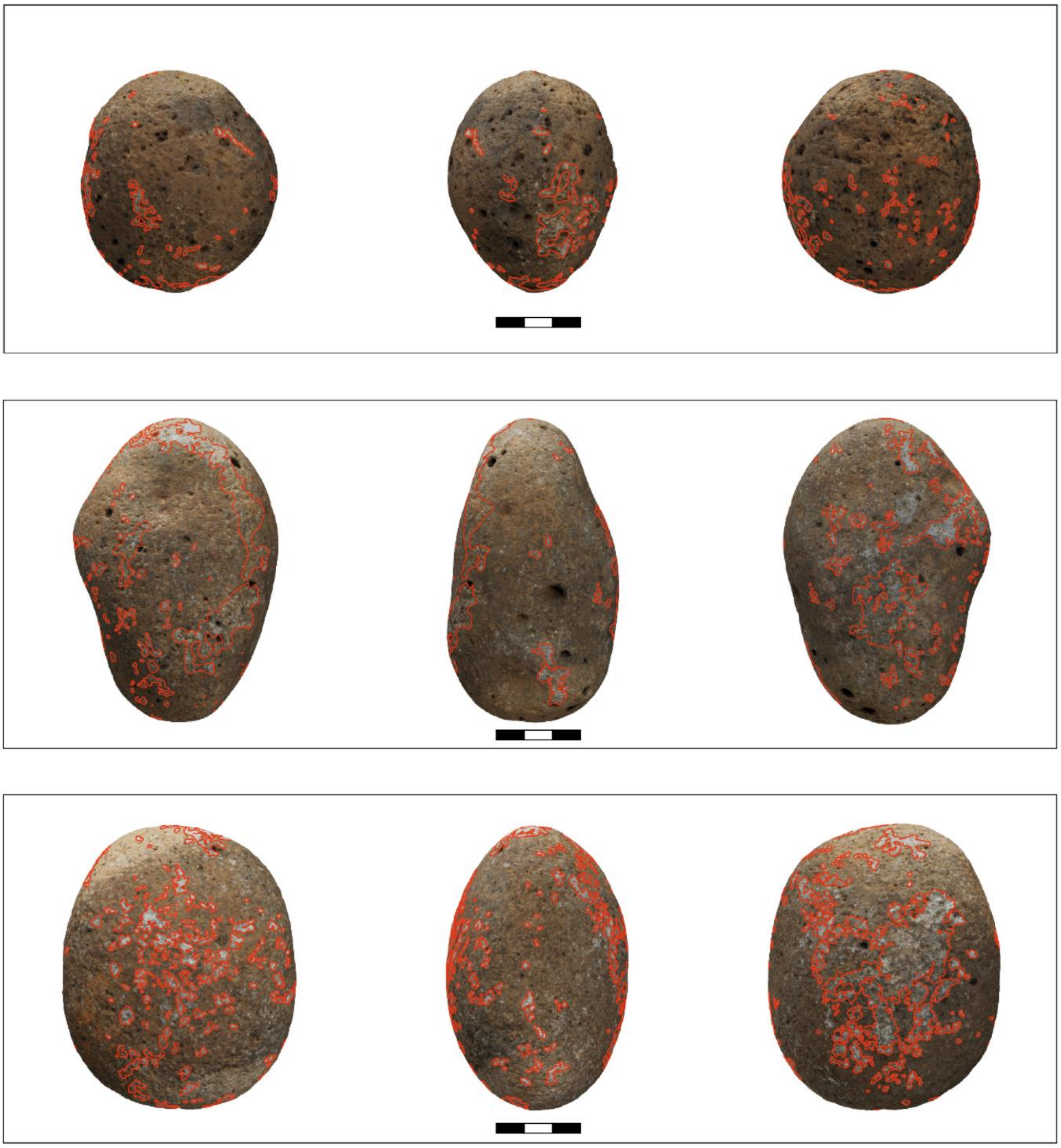

**Supplementary Figure 2:**
Use wear on capuchin hammerstones. Left to right: Plane A1, plane B1, and plan A2 of each stone. Small, medium and large hammer (top to bottom). Grey square: Scale of 10mm.

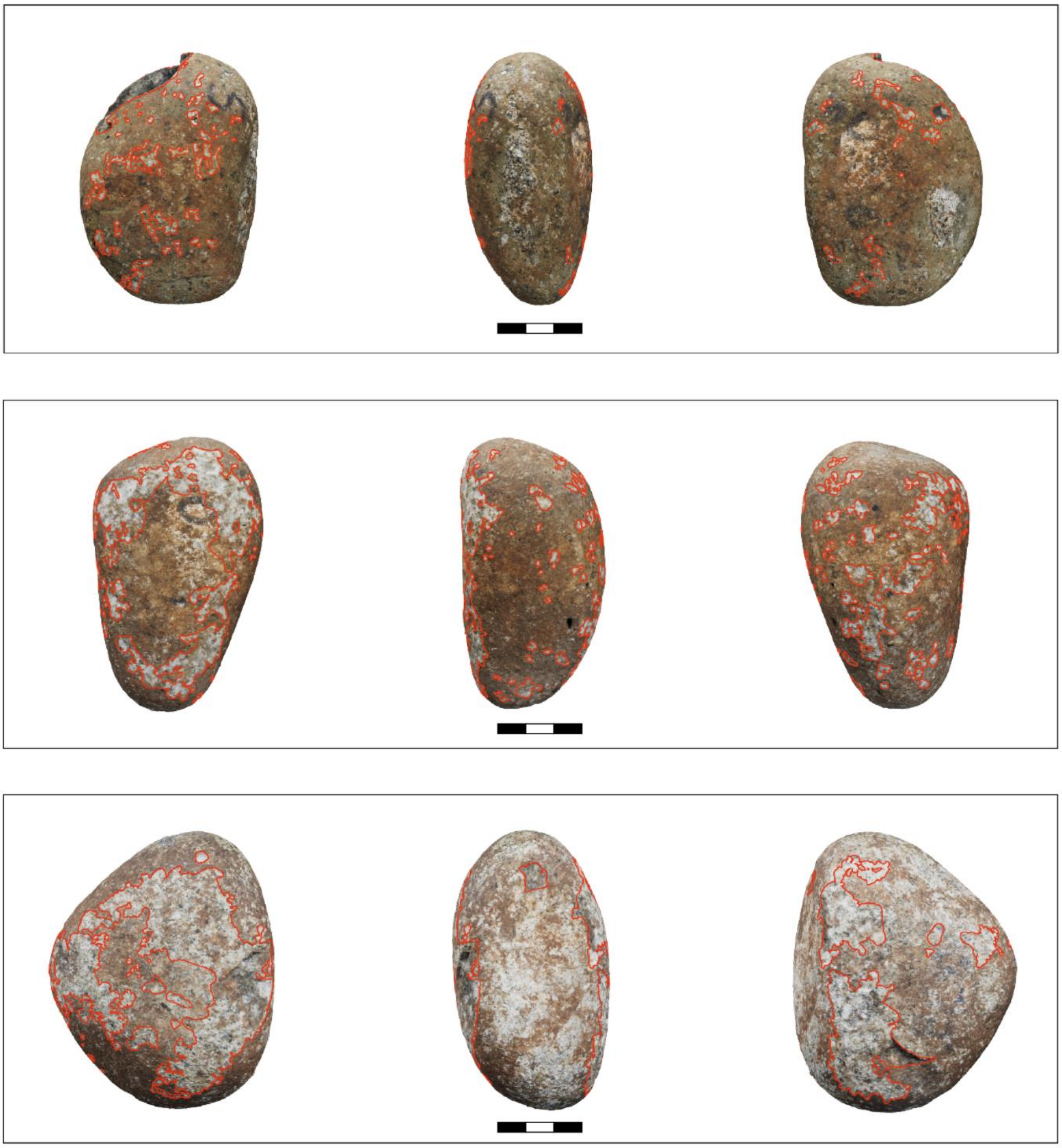

**Supplementary Figure 3:**
Use wear on macaque hammerstones. Left to right: Plane A1, plane B1, and plan A2 of each stone. Small, medium and large hammer (top to bottom). Grey square: Scale of 10mm.

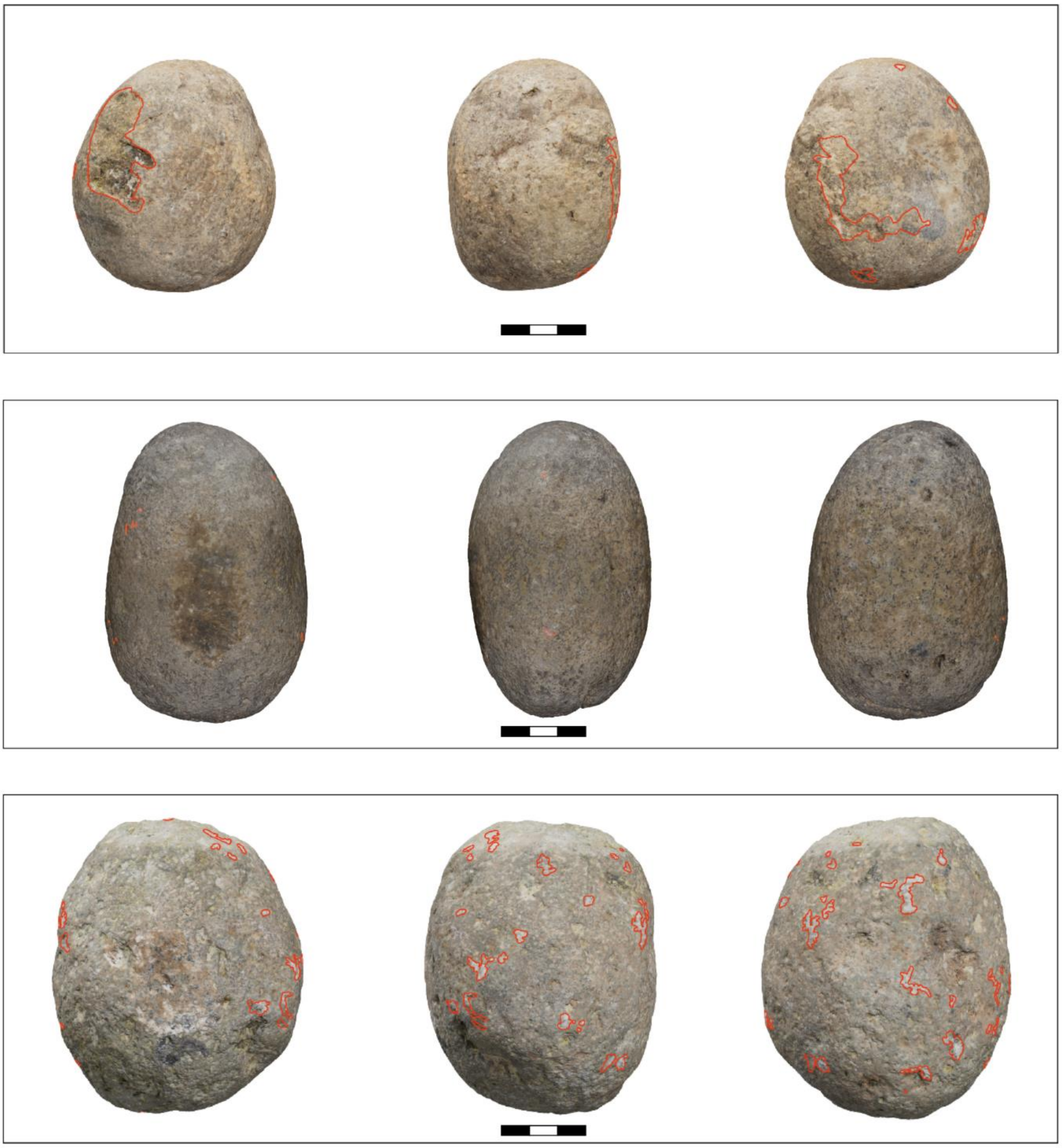

**Supplementary Table 20**: Use wear details of each hammerstone (“Small”, “Medium”, “Large”) of each species. Quantity of direct use wear areas; maximum, minimum, mean, and SD of use wear areas; total area covered in use wear; percentage of surface area covered in use wear (PA); largest use wear area (LUW); density of use wear areas; edge density of use wear areas (ED).

| **Use Wear** | **Chimpanzee** | | | | **Capuchin** | | | | **Macaque** | | | |
|---|---|---|---|---|---|---|---|---|---|---|---|---|
| | **Small** | **Medium** | **Large** | **Mean** | **Small** | **Medium** | **Large** | **Mean** | **Small** | **Medium** | **Large** | **Mean** |
| Num discreet use wear areas | 120 | 120 | 284 | 174.67 | 67 | 94 | 33 | 64.67 | 6 | 8 | 53 | 22.33 |
| Min Area (mm2) | 0.09 | 0.02 | 0.05 | 0.05 | 0.06 | 0.02 | 0.34 | 0.14 | 5.99 | 0.48 | 0.28 | 2.25 |
| Max Area (mm2) | 261.56 | 489.65 | 906.01 | 552.40 | 56.55 | 1016.42 | 2864.94 | 1312.64 | 392.16 | 5.51 | 42.33 | 146.67 |
| Mean Area (mm2) | 7.53 | 16.19 | 8.58 | 10.77 | 7.62 | 28.84 | 142.28 | 59.58 | 91.56 | 2.16 | 10.51 | 34.74 |
| SD Area (mm2) | 28.05 | 61.83 | 55.18 | 48.35 | 12.46 | 116.79 | 507.04 | 212.10 | 138.60 | 1.73 | 11.11 | 50.48 |
| Total Area (mm2) | 903.94 | 1942.38 | 2437.3 | 1761.21 | 510.34 | 2711.36 | 4695.22 | 2638.97 | 549.35 | 17.32 | 556.87 | 374.51 |
| PA (%) | 6.02 | 9.94 | 10.94 | 8.97 | 3.69 | 17.60 | 25.17 | 15.48 | 3.33 | 0.08 | 1.96 | 1.79 |
| LUW (%) | 1.74 | 2.50 | 4.07 | 2.77 | 0.41 | 6.60 | 15.36 | 7.46 | 2.38 | 0.03 | 0.15 | 0.85 |
| Density | 0.008 | 0.0061 | 0.0128 | 0.009 | 0.0048 | 0.0061 | 0.0018 | 0.0042 | 0.0004 | 0.0004 | 0.0019 | 0.0009 |
| Mean Density | 0.0005 | 0.0008 | 0.0004 | 0.0006 | 0.0006 | 0.0019 | 0.0076 | 0.0034 | 0.0055 | 0.0001 | 0.0004 | 0.002 |
| ED | 0.12 | 0.14 | 0.24 | 0.17 | 0.09 | 0.18 | 0.13 | 0.13 | 0.02 | 0.003 | 0.04 | 0.02 |

**Supplemantary Table 21**: Damage pattern of anvils across three primate species.

| | **Capuchin** | **Chimpanzee** | **Macaque** |
|---|---|---|---|
| **Tool** | | | |
| Active plane area ($cm^2$) | 221.44 | 215.48 | 261.40 |
| **Use Wear** | | | |
| Max Area ($cm^2$) | 83.32 | 0.9005 | 0.0235 |
| Min Area ($cm^2$) | 0.020 | 0.0170 | 0.0078 |
| Mean Area ($cm^2$) | 2.000 | 0.1291 | 0.0147 |
| SD Area ($cm^2$) | 11.42 | 0.2367 | 0.0063 |
| Total Area ($cm^2$) | 103.98 | 1.5490 | 0.0588 |
| PA (%) | 46.96 | 0.72 | 0.02 |
| LUW (%) | 37.63 | 0.42 | 0.009 |
